# Fluorescent-based sex-separation technique in major invasive crop pest, *Drosophila suzukii*

**DOI:** 10.1101/2024.10.07.617099

**Authors:** Junru Liu, Danny Rayes, Minzhe Yang, Omar S. Akbari

**Author notes:** To whom correspondence should be addressed: Omar S. Akbari, Ph.D., School of Biological Sciences, Department of Cell and Developmental Biology, University of California, San Diego, La Jolla, CA 92093, USA. equal contributions (Junru Liu, Danny Rayes).

## Abstract

Insect population biocontrol methods such as the sterile insect technique (SIT), represent promising alternatives to traditional pesticide-based control applications. To use these strategies efficiently requires scalable sex separation techniques which are currently lacking in *Drosophila suzukii*, a prominent crop pest species. Having previously characterized a fluorescence-based sex-sorting technique in other pests, termed SEPARATOR (<u>S</u>exing <u>E</u>lement <u>P</u>roduced by <u>A</u>lternative <u>R</u>NA-splicing of <u>A T</u>ransgenic <u>O</u>bservable <u>R</u>eporter), here we explore its potential applicability to *Drosophila suzukii*. Here, we engineer several strains of *Drosophila suzukii* encoding SEPARATOR constructs that allow for efficient sex selection in early larval stages.

## Introduction

The Spotted Wing Drosophila (SWD), *Drosophila suzukii (D. suzukii)*, has recently emerged as a major agricultural pest, rapidly spreading from East Asia to North America and Europe.^1^ Unlike most of its close relative species, *D. suzukii* oviposits eggs inside ripening fruit rather than decaying fruit.^2–4^ The females use a serrated ovipositor to pierce fruit and deposit eggs, resulting in larvae consuming the fruit.^2,5^ Additionally, the physical damage from oviposition creates entry points for secondary infections by bacteria, fungi, and yeast.^6,7^ Moreover, *D. suzukii’s* short generation time accelerates its spread, leading to significant economic losses and increased pesticide usage, including broadspectrum organophosphates, pyrethroids, and spinosyns.^8–11^

Alternative biocontrol methods, such as the sterile insect technique (SIT) offer sustainable alternatives to pesticides. These methods require mass-rearing of insects that are sterilized and released to suppress wild populations.^12^ In traditional SIT programs, such as those targeting the New World Screwworm, both sterile males and females are released together due to the lack of sex-sorting technologies.^13,14^ However, releasing only males has proven more cost-effective and efficient.^15^ Moreover, sterile females can damage crops through oviposition and reduce suppression efficiency by mating with sterile males making removal of females prior to release attractive.^16–18^

Currently, sex sorting is mainly done manually, which is labor-intensive, slow, and prone to human error. ^13,19^ Although machine learning has been applied to automate sex soring in mosquito species like *Aedes albopictus* and *Aedes aegypti*, these methods rely on natural sexual dimorphisms, limiting their use in other pests/vector species.^13,19,20^ The large-scale implementation of SIT requires an accurate, efficient, and adaptable alternative to these manual methods. Recently, pgSIT was developed in *D. suzukii*, eliminating the need for irradiation to produce sterile males and remove females.^21^ However, it still requires a manual setup of crosses to generate the released F1 sterile males. Combining pgSIT with SEPARATOR could potentially reduce the need for manual sorting and improve overall production efficiency and scalability.

In some species, genetic Sexing Strains (GSS) have been developed to enable sex sorting based on traits like temperature sensitivity, chemical sensitivity, or pupal coloration.^15,19,22–27^ For instance, pupal color has been used successfully in Mediterranean fruit flies (*Ceratitis capitata*, Medfly), where a Y-linked translocation enables efficient separation of females using sorting machines.^28–30^ Engineering a Y-linked GSS requires translocating a conditional lethal gene such as resistance to the insecticide dieldrin (Rdl) to the Y chromosome, providing resistance only to males.^31,32^ While Y-linked GSS can work, limitations such as partial sterility and genetic instability due to recombination and phenotype loss complicate their use.^27,33^ Therefore, a more flexible sorting mechanism applicable to various species is desirable. In response to the limitations of Y-linked GSS, autosomal linked GSS have been developed in several species.^34–36^ Many of these new systems leverage sex-specific alternative splicing of the conserved sex determination gene *transformer* (*tra*).^34,36–39^ In this study, we demonstrate that the Sexing Elements Produced by Alternative RNA-splicing of a Transgenic Observable Reporter (SEPARATOR), previously characterized in *Drosophila melanogaster, Ceratitis capitata*, and *Aedes aegypti*, also functions in *D. suzukii*.^40–42^ SEPARATOR uses alternative splicing, typically a posttranscriptional process that increases protein diversity, to allow sex-specific expression of a fluorescent marker. Our expression cassette contains a fluorescent protein with a sex-specific intron, enabling only one sex to express fluorescence after sex-specific RNA splicing. We used the female-specific *transformer* intron (*traF*) from *D. suzukii, D. melanogaster*, and *C. capitata* to demonstrate the system’s feasibility in *D. suzukii*.^41^ SEPARATOR’s functionality in *D. suzukii* represents a promising tool for integrating existing biocontrol methods and automatic sex sorting technologies.

## Result

### SEPARATOR resulted in Female-specific dsRed in *D. suzukii*

We engineered the SEPARATOR cassettes for *D. suzukii* by incorporating the female-specific *transformer* (*traF*) intron from three species--*C. capitata (CctraF), D. melanogaster (DmtraF)*, and *D. suzukii (DstraF)--*to generate 795H, I, and J constructs (**Fig. 1A**). In these constructs, *traF* was inserted into the coding sequence of dsRed immediately downstream of the ATG start codon and expressed under the control of the *Opie2* promoter. Each construct also included an eGFP dominant marker driven by the Hr5ie1 promoter. Additionally, a control construct (795G) was designed to express both dsRed and eGFP in both sexes. Multiple transgenic lines were generated for each construct through *piggyBac* transformation, but only one line with the most robust *eGFP* expression was selected for further testing. In the strains harboring 795H, I, and J, we only observed dsRed+/eGFP+ females and dsRed-/eGFP+ males, indicating female-specific dsRed expression (**Fig.1B)**.

**Fig 1:**
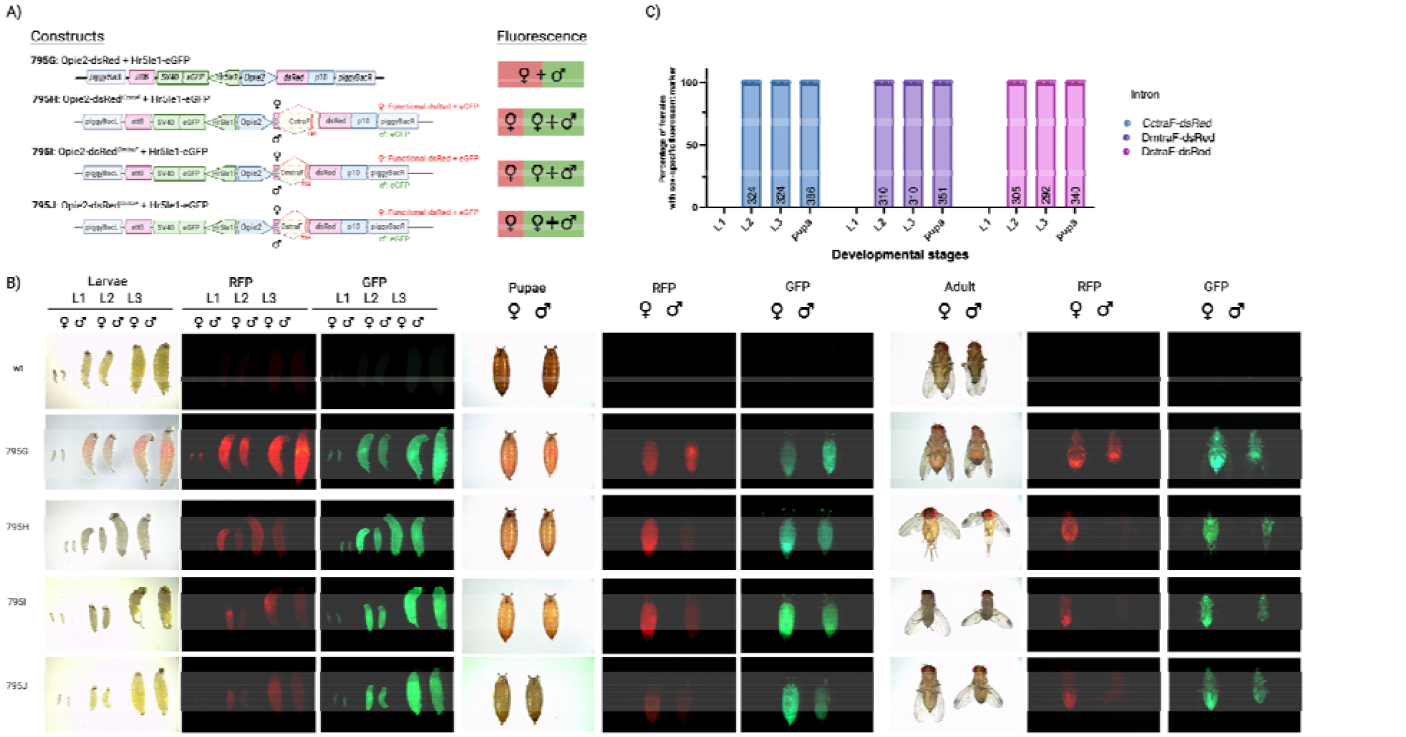
(**A**) Schematic of the SEPARATOR constructs developed and tested for the study, along with the corresponding fluorescence. (**B**) The expression of Opie2-*TraF*-dsRed in *D. suzukii* in L1-L3 larval, pupal, and adult stages for *CctraF, DmtraF*, and *DstraF* under white light, RFP, and GFP filters (**C**) Female selection efficiency at different developmental stages for all three constructs that exhibit female-specific dsRed. The number of scored flies is indicated in each bar. All three SEPARATOR constructs achieved 100% accuracy at sex separation; no males were detected with female-specific dsRed.

To verify female-specific splicing at the molecular level, pupae were collected for genomic DNA (gDNA) and RT-PCR analysis. The gDNA analysis showed identical band sizes in both females and males (**Fig. s1A**). In contrast, RT-PCR revealed multiple cDNA bands in both sexes (**Fig. s1B**). All females exhibit a 653 bp band, indicating *traF* splicing. In particular, in flies with *CctraF*, females show a single cDNA product, while males exhibit three cDNA products corresponding to three male-specific splicing patterns (**Fig. s1B**). In flies with *DmtraF*, females contain three cDNA products: one corresponding to the complete splicing, resulting in a 653bp dsRed product, and two additional products of 828bp and 901bp, corresponding to male-specific splicing and incomplete splicing, respectively (**Fig. s1B)**. In contrast, *DmtraF-dsRed* males only contain the 828bp and 901bp cDNA products, reflecting the male-specific splicing and incomplete splicing. Similarly, in flies with *DstraF*, females also contain three cDNA products: one for complete splicing (653 bp), and two for male-specific splicing (816 bp) and incomplete splicing (884 bp), respectively. Males with *DstraF*, like those with *DmtraF*, exhibit the 816 bp and 884 bp products, indicating male-specific splicing patterns and incomplete splicing. Sequencing results further confirmed the identity of these bands (**Fig. s1C & D**). All three constructs harboring *DmtraF, DstraF*, and *CctraF* introns resulted in female-specific dsRed in *D. suzukii* (**Fig. 1B**), consistent with the results observed in *D. melanogaster*.^*41*^

### SEPARATOR enables sex sorting in early larva

We evaluated the sex-specific fluorescence and identified the second instar larval (L2) stage as the earliest stage at which female-specific dsRed expression could be detected in the SEPARATOR strains (**Fig. 1C**). The eGFP fluorescence was similar across the four homozygous strains throughout the entire life developmental stages (795G, H, I, and J). However, the dsRed signal was notably more intense in females with *CctraF* compared to those with *DmtraF* and *DstraF*, despite *CctraF* being a non-endogenous intron. This stronger dsRed expression with *CctraF* persisted from the L2 stage through to the adult stage (**Fig. 1B)**.

### SEPARATOR Strains Maintain Fitness Comparable to Wild Type

Fitness is crucial for mass-rearing, as unfit flies can lower rearing efficiency. To determine whether SEPARATOR cassettes impact fitness, we measured egg hatching and larval-to-adult survival rates in three strains containing the *CctraF, DmtraF*, and *DstraF* (795H, I, and J ), comparing them to a constitutively expressed line (795G) and wild-type flies (**Fig.s2A & B, Table S2**). No significant differences in embryo hatching or larval-to-adult survival rates were observed between the transgenic strains, 795G, and the wild type (p<0.05, One-Way ANOVA multiple comparisons), indicating that incorporating *traF* does not adversely affect fitness and that our SEPARATOR strains perform similarly to the wild type under the tested conditions.

## Discussion

In this study, we demonstrated the effectiveness of SEPARATOR, in the agricultural pest *Drosophila suzukii*. SEPARATOR uses sex-specific alternative splicing of the conserved *transformer* gene to allow female-specific expression of dsRed. Males and females constitutively express eGFP, but only females fully splice the *traF* intron to restore functional dsRed translation. We tested *traF* introns from three species—*CctraF, DmtraF*, and *DstraF*—in *D. suzukii*, achieving 100% accuracy. We opted to employ *piggyBac*-mediated transformation in the creation of lines due to its ubiquitous functionality across both model and non-model insect species.^44^ Several strains per construct were generated to compensate for positional effects on expression. Although only three strains were further tested, all resulted in female-specific *dsRed* expression and exhibited fluorescence from the L2 instar onward.

RT-PCR confirmed that *CctraF, DmtraF*, and *DstraF* were spliced out completely in *D. suzukii* females, consistent with our observations in *D. melanogaster*. ^*41*^ Despite the random insertion by *piggyBac, CctraF* resulted in the strongest *dsRed* expression in *D. suzukii*. Specifically, only one cDNA product was observed for *CctraF* in *D. suzukii*, indicating 100% splicing efficiency and thus leading to more functional dsRed protein compared to *DmtraF* and *DstraF*. Moreover, *Cctra* does not require the presence of *sex-lethal* (*sxl)* in Tephritidae as opposed to Drosophilidae, which depends on *sxl* upstream for successful sex determination and, therefore, can possibly result in stronger expression.^45–47^ In *Drosophila*, sex is determined by the ratio of sex chromosomes to autosomes.^48,49^ An X ratio of 1 triggers the production of the sex-lethal (*sxl*) protein, which leads to the production of functional female-specific *transformer* (*tra*) protein. In contrast, in *Ceratitis, tra* is continuously autoregulated in XX females and suppressed in males by the maleness-on-the-Y (*MoY*) gene allowing reliable and stable splicing throughout the life cycle.^50,51,52^ We had previously predicted that *CctraF* could be a versatile tool for sex-specific expression across a wide range of insect species, and our findings in *D. suzukii* also support this conclusion. It should also be noted that *DmtraF* and *DstraF* should be employed primarily in species utilizing *sxl* as a master gene for somatic sex determination. Alternative splicing of *tra* in Drosophilidae relies on *sxl* for proper female splicing patterns.^47,53,54^ It’s also important to note that three bands were observed in *D*.*suzukii* males with *CctraF*, corresponding to the three male-specific transcripts (**Fig s1A & s1D**). This contrasts with the two male-specific transcripts reported in the endogenous *Ceratitis capitata* flies (**Fig. S3**).^34,42,55^

Nevertheless, we observed differences in *Cctra* splicing between males and females, suggesting the potential for male-specific splicing using *Ceratitis capitata transformer* male intron (*CctraM)* in *D. suzukii*. However, incomplete splicing of the *traF* intron or male-specific splicing of *tra* was observed in both *Dmtra* and *Dstra* in *D. suzukii*, indicating that male splicing patterns also occur in females. Thus, using the *D. melanogaster* and *D. suzukii transformer* male intron (*traM*) for male-specific expression in *D*.*suzukii* may not be feasible. Furthermore, fluorescent intensity varied widely between lines containing different introns which may be partially owed to the positional effect of the insertion site. Similarly, we also observed varying fluorescent intensity when characterizing SEPARATOR in *D. melanogaster* when all introns were inserted into the same site, indicating other factors were also at play. We attribute these differences to factors such as intron size, random insertion site, and variations in sex-determination pathways between *Ceratitis* and *Drosophila. CctraF* produced the brightest dsRed expression, followed by *DmtraF* and *DstraF*, a result consistent with our previous findings in *D. melanogaster*.^41^

One of SEPARATOR’s key advantages is its transferability across multiple species, as demonstrated in this study and previous work. Specifically, construct 795H1 (*CctraF*) enables successful sex-sorting in *D. melanogaster* and *C. capitata*, and in this study, we show that it achieves 100% accuracy in *D. suzukii*.^*41,42*^ This applicability is due to SEPARATOR’s use of *transformer*, a highly conserved gene in the sex determination pathway of many Dipteran species, allowing us to expand SEPARATOR into three different species in two genera with no modifications to our cassette and strategy.^40,41,56–58^

In combination with existing biocontrol methods, SEPARATOR can potentially be utilized in conjunction with a Complex Parametric Analyzer and Sorter (COPAS, Union Biometrica) instrument. COPAS has already proven successful in sorting *D. melanogaster* embryos and larvae, as well as *Aedes aegypti* and *Anopheles gambiae* larvae using fluorescence.^13,59^ The COPAS machine fully automated the sorting process, allowing efficiency, speed, and precision impossible in manual sorting.^13^

As previously mentioned, SEPARATOR may be susceptible to random genetic events resulting in loss-of-function, gain-of-function, or translocation and recombination, leading to loss of fluorescence.^41^ However, loss of fluorescence may not be remarkably detrimental to our system, as any mutations or events leading to loss of fluorescence will also be selected against in sorting. Even so, precaution and wariness to such events should be exercised, particularly at larger scales. Safeguards utilized in classic GSSs, such as the Filter Rearing System (FRS), should be implemented to protect against recombination.^60–62^ The FRS was explicitly designed for the Mediterranean fruit fly GSS program and functions by maintaining a small, regularly sorted colony on the side that can be quickly amplified and replaced in case of system breakdown.

With that, SEPARATOR could be applied in large-scale, automated sex sorting systems. Systems like incompatible insect technique (IIT) could benefit from such a highly accurate, automatable method due to the stringent need for male-only releases.^63,64^ SEPARATOR’s ability to sort as early as the L2 instar also represents significant cost savings for programs like SIT, which do not require sex sorting to function. Releasing sterile males and females in SIT still suppresses populations, but removing sterile females early reduces rearing costs, improves suppression efficiency by preventing sterile females from mating with sterile males, and is crucial when targeting disease vectors where females are responsible for transmission.^13,14,65–67^ Since our SEPARATOR system only contains the *tra* intron; it can be combined with systems (e.g. pgSIT) that use gRNA targeting the *tra* gene. As in those cases, the gRNA would specifically target the exon regions, thereby avoiding interference with the intron region utilized by SEPARATOR.^21^ Moreover, SEPARATOR does not rely on antibiotics for line maintenance or sex selection, further bolstering its cost-efficiency.^34,36,37,68^

## Methods

### Molecular constructs design and assembly

We used the previously described plasmids harboring various *traF* introns inside the dsRed coding sequence and the *Hr5ie1-eGFP* markers to generate the SEPARATOR strains: *Opie2-dsRed, Opie2-CctraF-dsRed, Opie2-DmtraF- dsRed*, and *Opie2-DstraF-dsRed* (Addgene ##205481-205484).^41^ All plasmids’ *traF* introns were PCR amplified from the genomic DNA of their respective species using primers listed in **Table S1**. Restriction endonucleases AvrII and BamHI were used to linearize the plasmids before insertion of the *traF* intron into the coding sequence of *dsRed* immediately downstream of the ATG start codon.

### Rearing and fly transgenesis

Transgenic flies were maintained under standard conditions at 21ºC with a 12H/12H light/dark cycle and were fed on a cornmeal-yeast-agar diet in an ACL2 insectary at UCSD. Embryonic injections were performed following standard injection protocols in the lab. Plasmids were diluted to 300-350 ng/µL in water and then injected into the *D. suzukii* embryos that harbored the Hsp70Bb-piggyBacTransposase.^69^ G0 adults emerged from the injection were outcrossed to *D. suzukii wt* flies.

### Reverse transcription PCR (RT-PCR) of the female-specific splicing transcripts

To analyze the splicing transcripts of the three *traF* introns, dsRNA mRNA specific for females and males was screened. Total RNA was extracted from ten female or male pupae from wild-type 795G, H, I, and J strains using the miRNeasy Tissue/Cells Advanced Kits (Qiagen). Subsequently, DNase treatment was performed using the TURBO™ DNA-free kit (Invitrogen). The cDNA synthesis was conducted with the RevertAid First Strand cDNA Synthesis Kit (Thermo Scientific™). Genomic DNA (gDNA) was amplified using 795.s2F and 795.s2R, while cDNA was amplified using primers 795.s3F and 795.s1R (Supplementary Table 1). The gDNA samples were analyzed on a 1% TAE agarose gel, and the cDNA samples were analyzed on a 2% TAE agarose gel.

### Genetics and sex selection

To evaluate the effectiveness of the fluorescent sex selection, we paired ten virgin females with ten males in a fly embryo collection chamber. New grape plates were replaced every 12 hours, and the numbers of the laid embryos were recorded and transferred to fresh fly vials. After hatching, the larvae or pupae were sorted into vials based on their fluorescent markers. The sex and fluorescent markers of the adult offspring were documented after eclosion. Flies were examined using a Leica M165FC fluorescent stereomicroscope, and images were captured using a View4K camera. Each genetic cross was done four times with different parental flies.

### Fitness estimation

Two parameters were used to assess the fitness of the sex-sorting strains: the egg-hatching rate (from embryos to larvae), and the adult survival rate (from larvae to adult stage). To measure the egg-hatching rate, flies were allowed to lay embryos in the fly embryo collection chamber for 24 hours, embryos were collected and placed onto black filter paper atop the larval diet. The number of unhatched eggs were recorded after 24 hours and subtracted from the total to achieve the hatching rate. The number of female and male adult flies that successfully emerged is recorded to evaluate the adult survival rate.

### Statistical analysis

Statistical analysis was performed in Prism9 by GraphPad Software, LLC. Three to five biological replicates were used to generate statistical means for comparisons.

## Data availability

Complete sequence maps and plasmids are deposited at Addgene.org (#205481-205484). Transgenic lines are available upon request to O.S.A.

## Acknowledgments

We would like to thank Annie Kim for help with fly work.

## Author contributions

O.S.A. and J.L conceived and designed the experiments. J.L, D.R, and M.Z performed molecular and genetic experiments. All authors contributed to the writing, analyzed the data, and approved the final manuscript.

---

**Fig s1.**
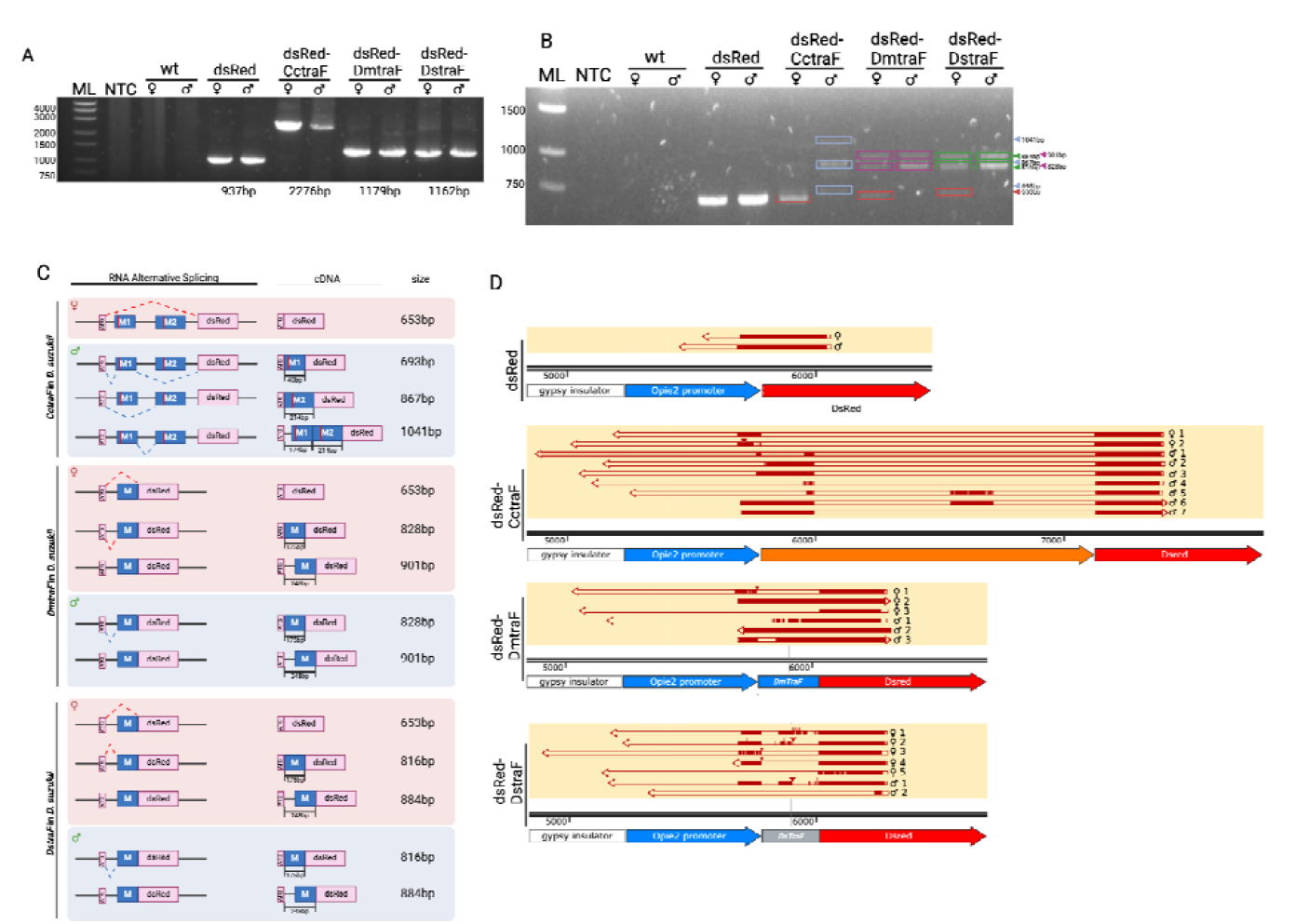
Gel electrophoresis images show (**A**) the genomic DNA PCR and (**B**) RT-PCR for *dsRed-traF*. (**C**) RNA alternative splicing and resulting cDNA, and (**D**) sequencing results of the RT-PCR of *CctraF, DmtraF*, and *DstraF*.

**Fig s2.**
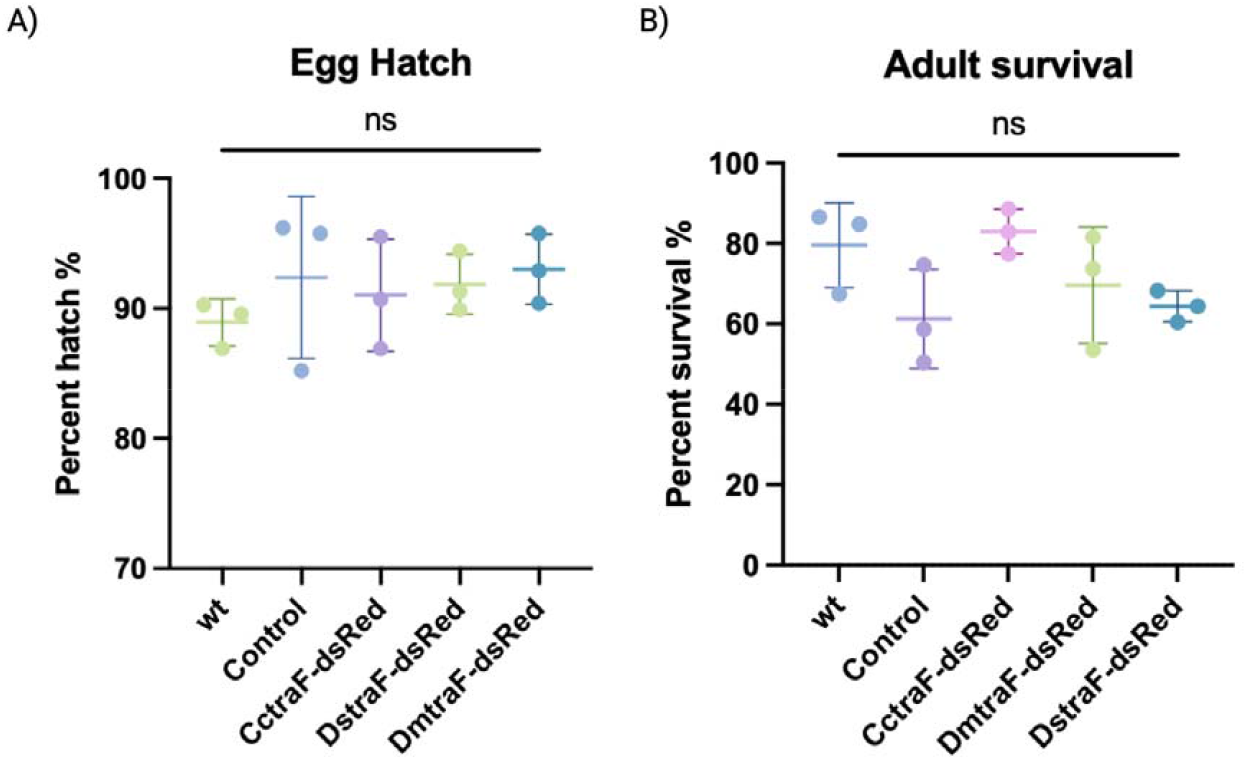
The fitness of the SEPARATOR is measured through (**A**) Embryo hatching rate and (**B**) survival rate from larvae to adults. No fitness cost was observed when comparing *wildtype* ( p<0.05, One-Way ANOVA multiple comparisons).

**Fig S3.**
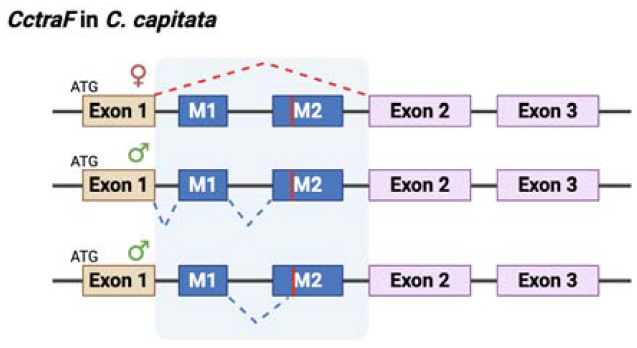
Alternative Splicing of *CctraF* in *C. capitata*.

**Table S1:**
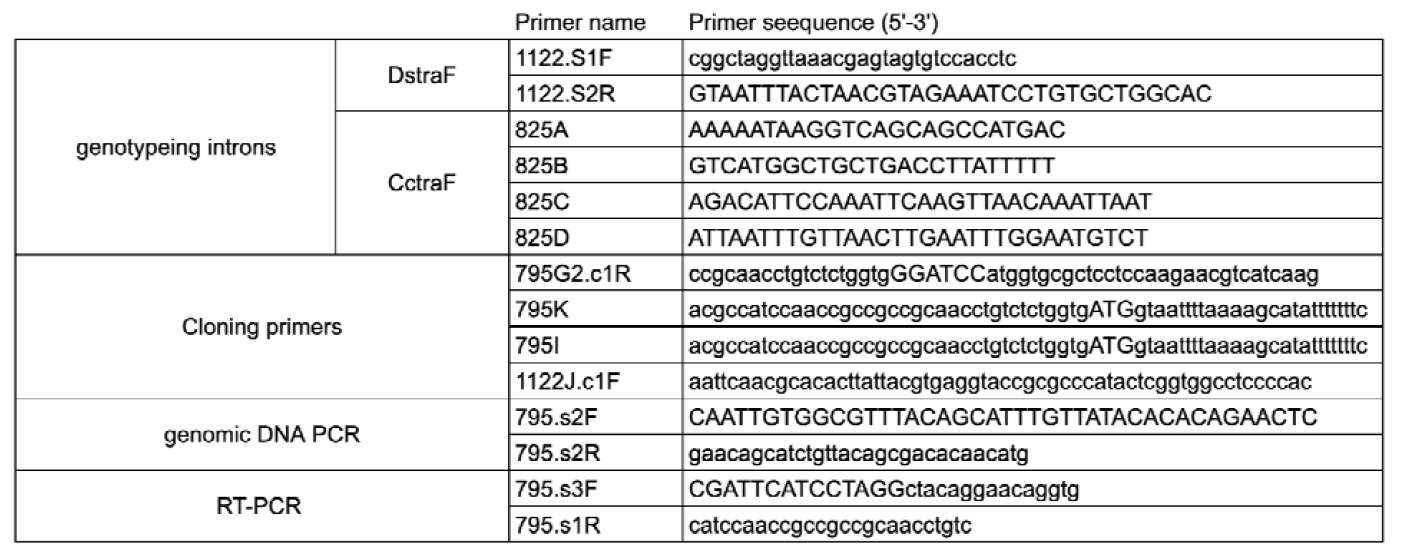
Primer sequences used in this study.

**Table S2:**
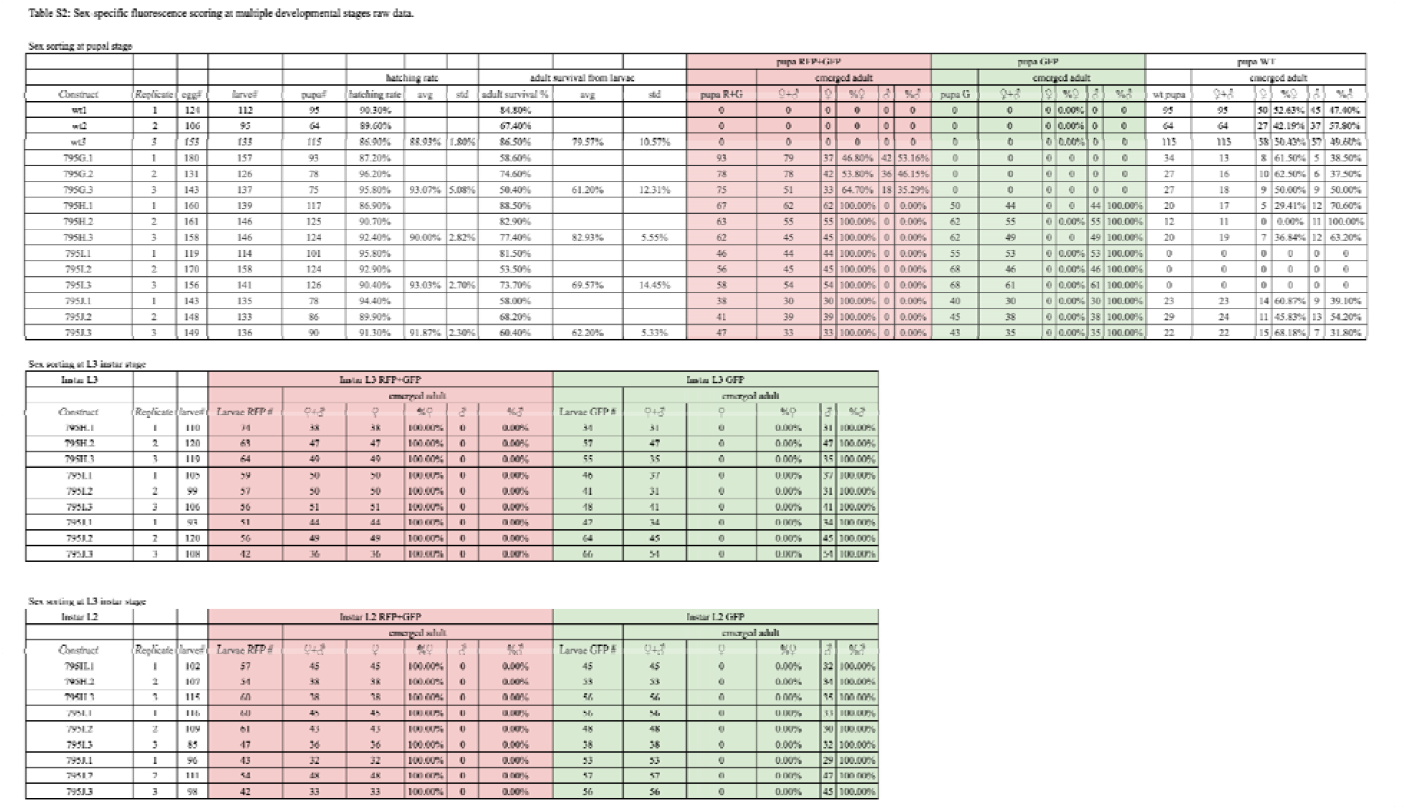
Sex-specific fluorescence scoring at multiple developmental stages raw data.

